# Intrinsic Buffer Hydroxyl Radical Dosimetry Using Tris(Hydroxymethyl)Aminomethane

**DOI:** 10.1101/825463

**Authors:** Addison E. Roush, Mohammad Riaz, Sandeep K. Misra, Scot R. Weinberger, Joshua S. Sharp

**Affiliations:** Department of BioMolecular Sciences, University of Mississippi, Oxford, Mississippi 38677, United States; Department of Chemistry and Biochemistry, University of Mississippi, Oxford, Mississippi 38677, United States; GenNext Technologies, Inc., Montara, California 94037, United States

## Abstract

Fast Photochemical Oxidation of Proteins (FPOP) is a powerful covalent labeling tool that uses hydroxyl radicals generated by laser flash photolysis of hydrogen peroxide to footprint protein surfaces. Because radical production varies with many experimental parameters, hydroxyl radical dosimeters have been introduced to track the effective radical dosage experienced by the protein analyte. FPOP experiments performed using adenine optical radical dosimetry containing protein in Tris buffer demonstrated unusual dosimetry behavior. We have investigated the behavior of Tris under oxidative conditions in detail. We find that Tris can act as a novel gain-of-signal optical hydroxyl radical dosimeter in FPOP experiments. This new dosimeter is also amenable to inline real-time monitoring thereby allowing real-time adjustments to compensate for differences in samples for their quenching ability.

## Introduction

Since its introduction by Hambly and Gross [1], Fast Photochemical Oxidation of Proteins (FPOP) coupled with mass spectrometry (MS)-based bottom-up proteomic methods has become a powerful technique for characterizing protein topography. FPOP relies on characterizing protein topography by measuring the apparent rate of reaction of amino acid side chains with diffuse hydroxyl radicals generated by laser flash photolysis of hydrogen peroxide. The apparent oxidation rate of each amino acid is dependent on both its inherent reactivity (which in turn is dependent on the sequence context [2, 3] and side chain structure [4]) and the radical accessibility of the side chain [3, 5-7]. These labeling reactions produce covalently bound, stable modification products which are unaffected by downstream sample-handling [8-10], and, though it currently provides lower resolution than other structural biology techniques such as X-ray crystallography, NMR, and cryo-EM, FPOP benefits from a low sample-size requirement, tremendous flexibility in sample characteristics (homogeneity, size, dynamics, etc.), and the ability to complete initial protein-radical chemistry on a low microsecond timescale that is faster than conformational changes can occur [1, 8, 9], although secondary reactions can persist longer [11, 12].

Structural characterization by FPOP typically depends on comparing protein footprints obtained under several conditions. However, alterations to hydroxyl radical scavenging capacity due to changes in buffer composition or the addition of some ligands and/or binding partners make it difficult to standardize results for comparison between experiments and between labs. To overcome this issue, several molecules have been introduced to the FPOP workflow which allow the effective hydroxyl radical concentration experienced by the analyte to be determined, thereby providing a metric with which experiments can be compared [13-16]. Each of these dosimeters competes with the analyte for hydroxyl radicals and experiences a change in its measurable properties proportional to the amount of radical present and not scavenged by other pathways.

The UV-absorbent molecule adenine offers an easy option for radical dosimetry [15, 17], and although it initially necessitated the introduction of additional steps to the FPOP workflow, the recent introduction of an inline UV-spectrometer [10] negates this issue and allows hydroxyl radical production to be monitored in real-time. Adjustments to peroxide concentration, laser fluence, and scavenging capacity can then be made as an experiment is performed to maintain a consistent level of oxidation across all samples [18]. Recently, while performing FPOP experiments in Tris buffer with the adenine dosimeter, we observed adenine dosimetry readings that were inconsistent with protein oxidation and exhibited unexpected gain of absorbance behavior. We investigated the properties of Tris under oxidative conditions and found that Tris acted as an effective intrinsic buffer/dosimeter that allows for optical hydroxyl radical dosimetry without the need to add an exogenous molecule; the buffer itself provides the hydroxyl radical dosimeter.

## Material and Methods

Tris, myoglobin, Glu1-fibrinopeptide B (GluB), and MES hydrate were purchased from Sigma-Aldrich Corporation (St. Louis, MO, USA). Hydrogen peroxide (30%) was purchased from J.T. Baker (Phillipsburg, NJ, USA), and sequencing grade-modified trypsin was purchased from Promega (Madison, WI, USA). Methionine amide was purchased from Bachem (Torrance, CA, USA). Dithiothreitol (DTT) was purchased from Soltec Ventures (Beverly, MA, USA). LC/MS-grade formic acid and LC/MS-grade acetonitrile were purchased from Fisher chemical (Fair Lawn, NJ, USA). 8.5 mM Tris was used in all experiments in order to maintain a hydroxyl radical scavenging capacity comparable to the 1 mM adenine and 17 mM glutamine used in many FPOP experiments [1, 9, 19]. All oxidation was performed by exposure to the pulsed beam of a COMPex Pro 102 KrF excimer laser (Coherent Inc., Santa Clara, CA) using standard FPOP methods described previously [1]. For offline dosimetry, ultraviolet absorbance was measured on a Thermo NanoDrop 2000c UV spectrophotometer. For real-time inline dosimetry, a Pioneer Series inline dosimeter (GenNext Technologies, Montara, CA) was used [10]. All samples were deposited after FPOP directly into a quench solution containing 0.5 µg/µL catalase and 0.5 µg/µL methionine amide to reduce secondary oxidation products.

Oxidized GluB and myoglobin samples were denatured by incubation at 90°C for 15 minutes, and digested by adding 1:20 w/w sequencing grade modified trypsin and incubating with rotation at 37 °C overnight. Digestion was stopped by adding 0.1% formic acid, and samples were analyzed by LC-MS/MS on an Orbitrap Fusion Tribrid mass spectrometer (Thermo Fisher Scientific). Myoglobin peptide separation was performed on an Acclaim PepMap 100 C18 nanocolumn (0.75 mm × 150 mm, 2 µm, Thermo Fisher Scientific). Peptides were eluted using a binary gradient of water with 0.1% formic acid (A) and acetonitrile with 0.1% formic acid (B) by increasing an initial 2% B gradient to 35% B over 22 minutes, ramping to 95% B over 5 minutes, holding for 3 minutes, returning to 2% B over 3 minutes, and finally holding at 2% B for 9 minutes. Electrospray voltage was set to 2500 V, and the ion transfer tube temperature was 300 °C. The top eight peaks from MS1 were fragmented by CID. Oxidation events per peptide were calculated using methods previously described [10].

## Results and Discussion

In order to investigate the behavior of Tris buffer under oxidative conditions, quadruplicate samples of Tris were reacted with hydroxyl radicals produced from the photolysis of hydrogen peroxide with an excimer laser. Upon comparing the UV absorbance spectra of Tris (obtained with a NanoDrop spectrometer) both with and without laser irradiation, we observed that, while Tris buffer is inherently absorbent in the short wavelength region of the ultraviolet spectrum, this absorbance changes little after oxidation by FPOP. In contrast, unoxidized Tris buffer was seen to be minimally absorbing in the longer wavelength UV region from 250-310 nm, but oxidation caused a substantial absorbance increase in this region with a maximum at 265 nm (**Figure S1**, Supplementary Information). In order to determine the source of Tris absorbance behavior, quadruplicate samples were oxidized under four different conditions as shown in **Figure 1**. As expected, all samples maintained a basal level of absorbance at 265 nm, but the absorbance significantly increased upon exposure to hydroxyl radicals. Based on these observations, we hypothesized that Tris buffer could be used as a hydroxyl radical dosimeter in FPOP reactions. Any such dosimeter must be capable of measuring the effective hydroxyl radical dose experienced by the protein analyte under diverse conditions. We performed FPOP on quadruplicate samples containing Tris and the model peptide GluB to evaluate the correlation between Tris absorbance and peptide oxidation. Laser fluence was held at 10.23 mJ/mm^2^ for all samples, and the concentration of hydrogen peroxide was varied from 5-40 mM in increments of 5 mM to generate increasing concentrations of radical. Because an increase in radical concentration corresponds to increased oxidation of analytes, we expected to see a direct, positive correlation between Tris absorbance and GluB oxidation. **Figure 2** shows that the absorbance of Tris at 265 nm does strongly positively correlate to average GluB oxidation per peptide, with deviations most likely dominated by error in the measurement of GluB oxidation [20].

**Figure 1.**
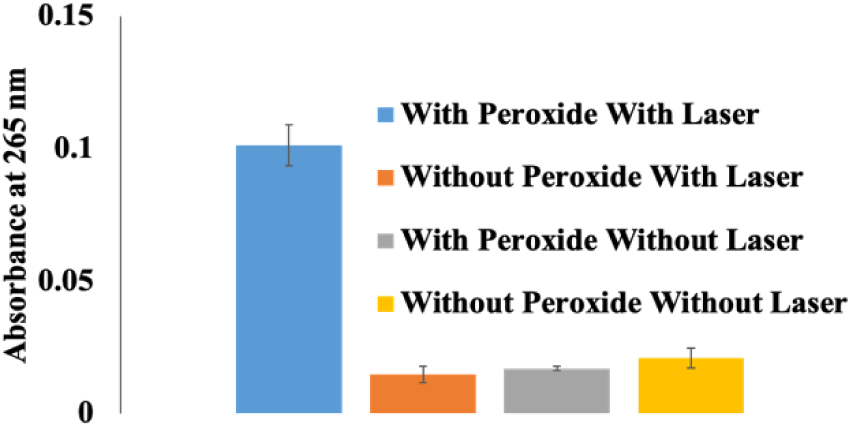
Absorbance of Tris at 265 nm as measured by NanoDrop after exposure to four sample conditions. Absorbance increases significantly only upon reaction with hydroxyl radicals generated by flash photolysis of hydrogen peroxide.

**Figure 2.**
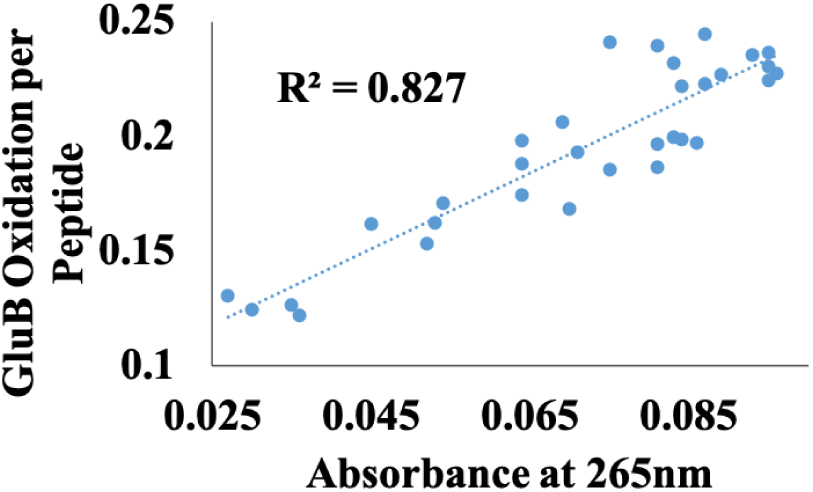
Tris absorbance at 265 nm correlates positively with average oxidation of GluB peptide.

We also observed that the gain in Tris absorbance was less when a competing radical scavenger, MES buffer, was added in the solution (**Figure S2**, Supplementary Information). Both of these observations support the hypothesis that Tris could be used as a potential dosimeter. However, it is important to note that both experiments used a simple sample mixture containing only Tris and one additional component. In most FPOP experiments, the sample contains several additional analytes, which compete with the dosimeter for radicals, so it is important to see that Tris maintains its dosimetry abilities under such complex conditions.

In order to test the robustness of Tris acting as a radical dosimeter, a standard FPOP reaction containing myoglobin was carried out with buffer pH held at 8.0 to maintain myoglobin conformational stability. The reaction mixture was oxidized in the presence as well as the absence of MES buffer, and absorbance readings were obtained in real-time using inline dosimeter monitoring. First, myoglobin was oxidized and the ΔAbs265 readings were 4.97±0.15 absorbance units, at a laser fluence of 11.66 mJ/mm^2^. In the presence of 10 mM MES buffer, the ΔAbs_265_ decreased to 3.37±0.30 absorbance units at 11.66 mJ/mm^2^, reflecting scavenging by the MES buffer. In a separate experiment, laser fluence was increased to 15.30 mJ/mm^2^ during the exposure of the myoglobin+MES sample to achieve a ΔAbs265 reading ≈ 4.97, identical to that of myoglobin without MES buffer (**Figure 3A)**. When FPOP is performed in the Tris buffer alone, the peptides are more oxidized; when MES is also added to the mixture, a drop in the oxidation of all myoglobin peptides is observed. By compensating for the scavenging capacity of MES buffer using Tris as a dosimeter, the compensated oxidation of all myoglobin peptides in the presence of MES buffer is the same as in the samples without MES scavenger as shown in **Figure 3B**, demonstrating that Tris can act as a functional and practical radical dosimeter for scavenging compensation [18].

**Figure 3.**
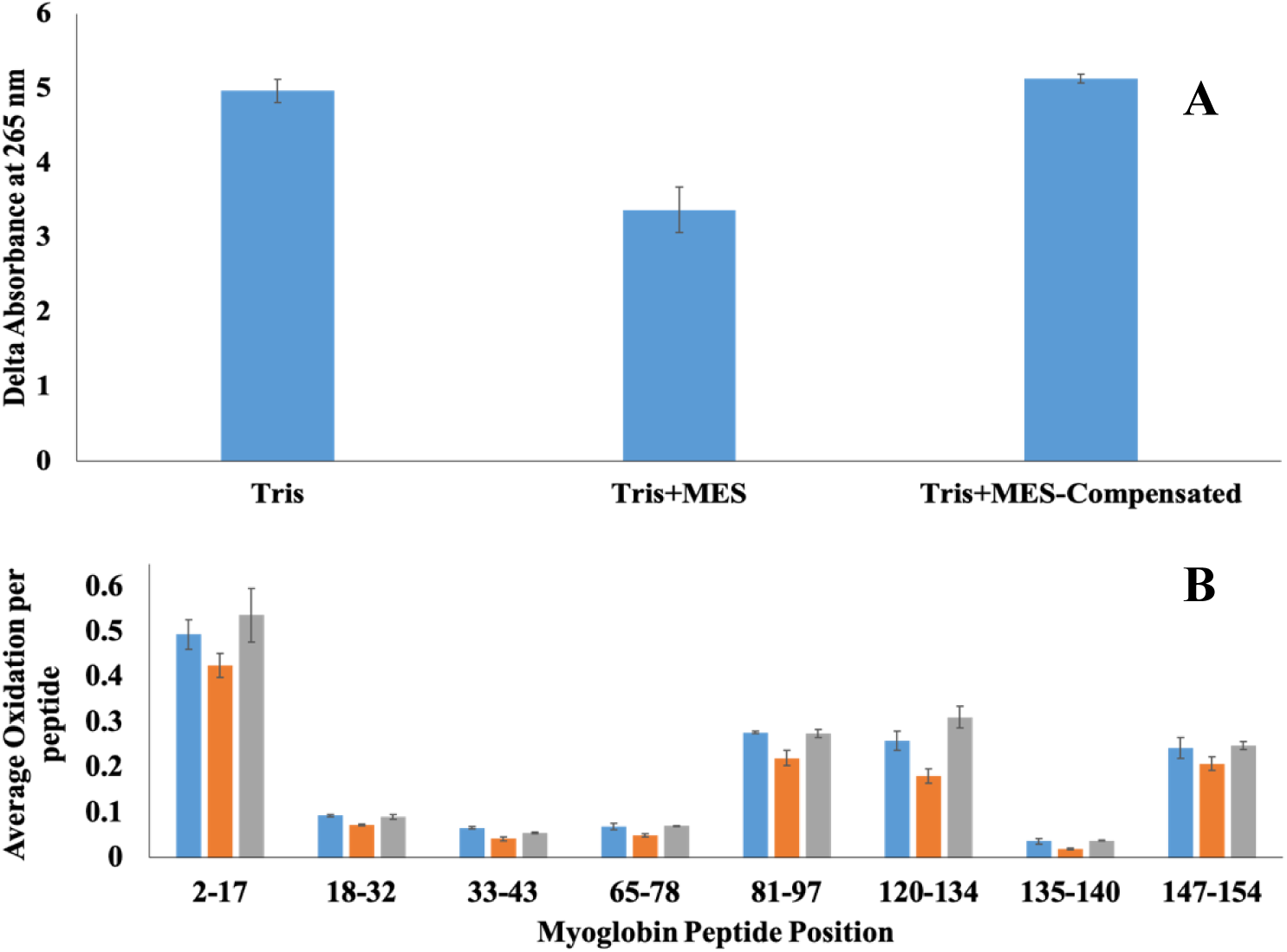
**A)** Tris absorbance change for myoglobin samples without MES scavenger, with 10 mM MES scavenger, and compensated conditions with 10 mM MES scavenger and increased laser fluence to obtain a ΔAbs_265_ ≈ 4.97. **B) (Blue)** Peptide oxidation for myoglobin peptides in the absence of MES; **(Orange)** Peptide oxidation for myoglobin peptides in the presence of 10 mM MES; **(Grey)** Peptide oxidation for myoglobin in the presence of 10 mM MES under compensating laser fluence conditions, using Tris as a doseimeter for radical compensation. No statistically significant differences were detected in peptide oxidation between no MES samples and with MES-containing samples compensated using Tris dosimetry.

## Conclusion

We have shown that the common buffer Tris(hydroxymethyl)aminomethane can be used as a highly effective hydroxyl radical dosimeter for FPOP experiments. Increases in Tris absorbance correlate strongly with peptide oxidation (R^2^ = 0.827) and scavenging capacity (R^2^ = 0.9625), and the absorbance loss resulting from increased scavenging capacity can be compensated in real-time to maintain consistent protein footprints. We suspect that this new chromophore is the result of formation of a *gem-*diol followed by water elimination, resulting in aldehyde and/or imine formation [21], with a proposed scheme as shown in **Figure S3**, Supplementary Information. Several characteristics of Tris suggest that it may be a favorable replacement for adenine dosimeter in many FPOP applications. Because the molecule is UV-active in the same range as adenine, no modifications to current measurement technologies are required for its adoption. As shown in **Figure S4**, Tris is the major contributor to absorbance change after laser exposure, so there is little interference from proteins or other buffer components. Tris also eliminates the need for the background scavenger glutamine thereby simplifying sample preparation. Furthermore, the use of Tris instead of adenine will allow for the application of FPOP to nucleoside and nucleotide binding proteins (a very large category of proteins) without concern about dosimeter interference in protein structure.

## Acknowledgments

This work was funded by the National Institute of General Medical Sciences (R01GM127267 and R43GM125420). We would like to thank Dr. Gerald B. Rowland and Dr. Daniell L. Mattern for many valuable discussions about possible reaction schemes resulting in Tris oxidation product absorbance behavior.

