## Supplementary Information for "Intrinsic Buffer Hydroxyl Radical Dosimetry Using Tris(Hydroxymethyl)Aminomethane"

### Supplemental Information

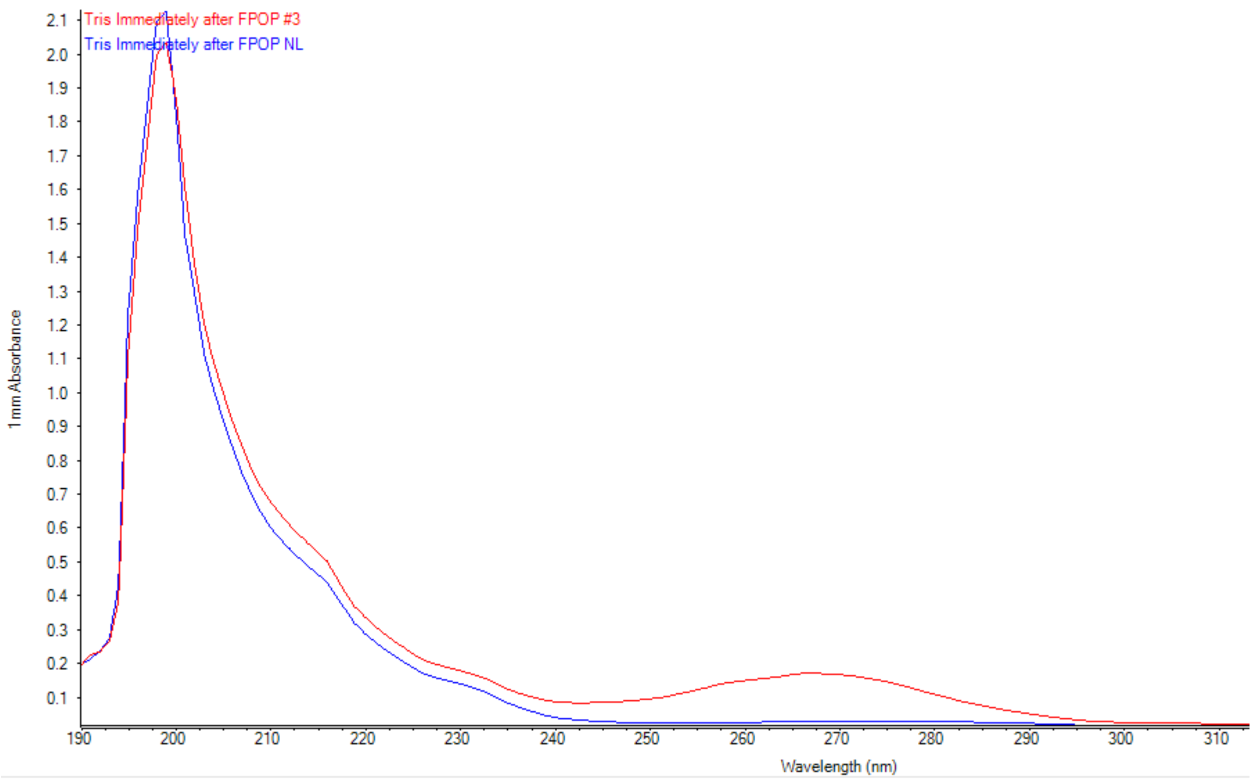


**Figure S1. Tris gains significant absorbance in the long wavelength UV region from 250-310 nm. Blue trace is 8.5 mM Tris with 100 mM peroxide. Red trace is 8.5 mM Tris with 100 mM peroxide and laser exposure.**

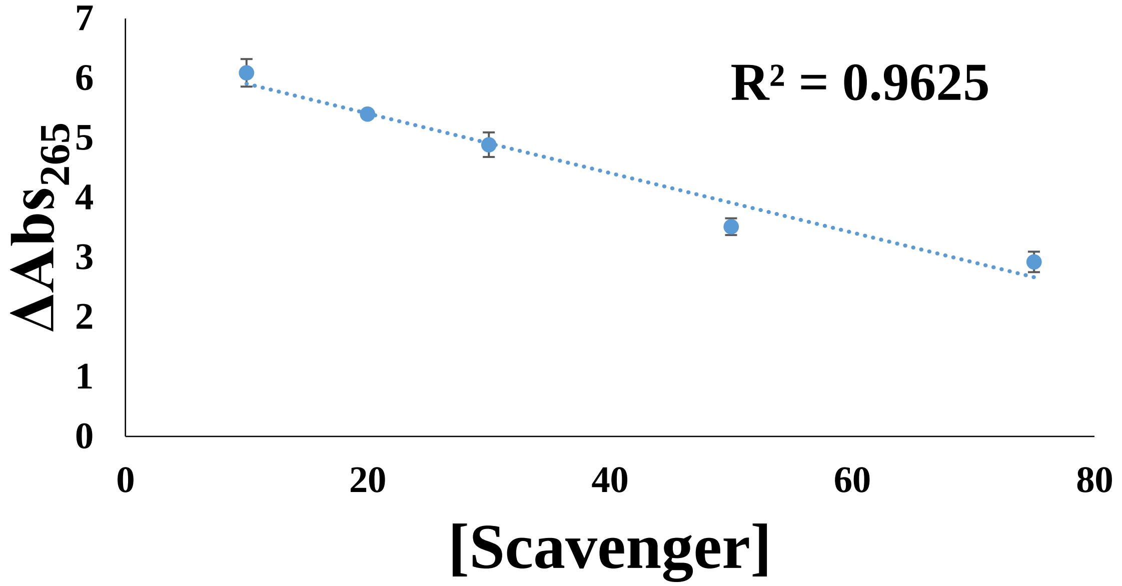


**Figure S2. The gain in Tris absorbance decreases with increasing MES concentration.**

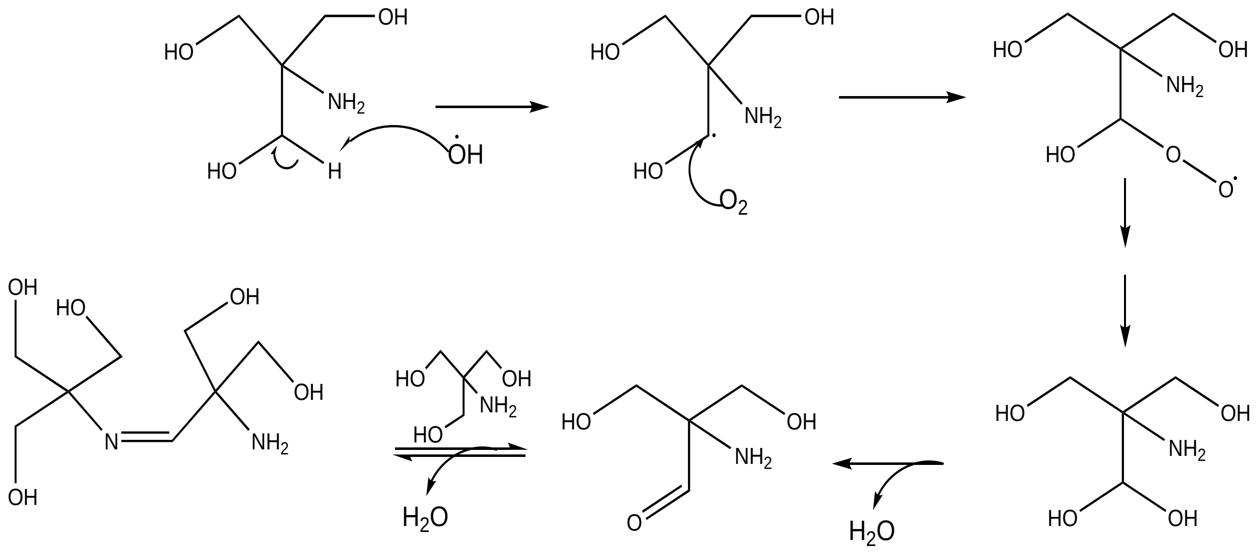


**Figure S3. Proposed scheme of Tris modification by hydroxyl radicals.**

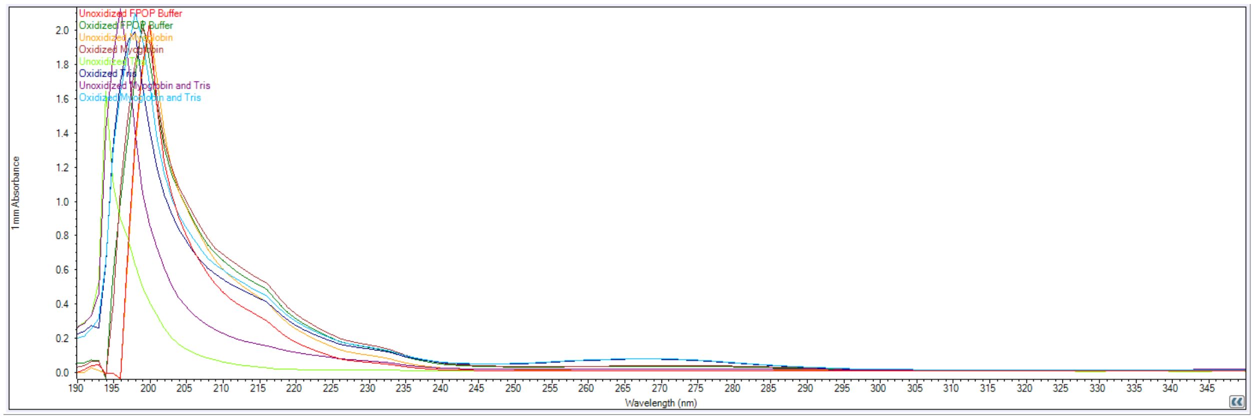


**Figure S4. UV absorbance spectra of FPOP sample components before and after oxidation by FPOP. pH was held at 8.01 for all samples, and 17 mM glutamine was used to maintain scavenging capacity in samples not containing Tris. Tris concentration was 8.5 mM, and myoglobin concentration was 5 µM. Oxidation was performed in 100 mM peroxide for all samples. Only Tris shows an appreciable change in UV absorbance at 265 nm.**
